# The ascending arousal system promotes optimal performance through meso-scale network integration in a visuospatial attentional task

**DOI:** 10.1101/2020.12.04.412551

**Authors:** Gabriel Wainstein, Daniel Rojas-Libano, Vicente Medel, Dag Alnæs, Knut K. Kolskår, Tor Endestad, Bruno Laeng, Tomas Ossandon, Nicolás Crossley, Elie Matar, James M. Shine

**Affiliations:** Brain and Mind Centre, The University of Sydney, Sydney, NSW, Australia; Centro de Neurociencia Humana y Neuropsicología, Facultad de Psicología, Universidad Diego Portales, Santiago, Chile; NORMENT, Division of Mental Health and Addiction, University of Oslo, and Oslo University Hospital, Oslo, Norway; Bjørknes University College, Oslo, Norway; Department of Psychology, University of Oslo, Oslo, Norway; Sunnaas Rehabilitation Hospital HT, Nesodden, Norway; RITMO Centre for Interdisciplinary Studies in Rhythm, Time and Motion, University of Oslo, Norway; Helgelandssykehuset Mosjøen, Helse Nord, Norway; Department of Psychiatry, School of Medicine, Pontificia Universidad Católica de Chile, Santiago, Chile; Institute for Biological and Medical Engineering, Schools of Engineering, Medicine and Biological Sciences, Pontificia Universidad Católica de Chile, Chile; Centre for Complexity, The University of Sydney, Sydney, NSW, Australia

**Author notes:** **Corresponding author**, James M. Shine –.

## Abstract

Previous research has shown that the autonomic nervous system provides essential constraints over ongoing cognitive function. However, there is currently a relative lack of direct empirical evidence for how this interaction manifests in the brain at the macro-scale level. Here, we examine the role of ascending arousal and attentional load on large-scale network dynamics by combining pupillometry, functional MRI and graph theoretical analysis to analyze data from a visual motion-tracking task with a parametric load manipulation. We found that attentional load effects were observable in measures of pupil diameter and in a set of brain regions that parametrically modulated their BOLD activity and meso-scale network-level integration. In addition, the regional patterns of network reconfiguration were correlated with the spatial distribution of the *α*2a adrenergic receptor. Our results further solidify the relationship between ascending noradrenergic activity, large-scale network integration, and cognitive task performance.

**Author Summary:** *In our daily lives, it is usual to encounter highly demanding cognitive tasks. They have been traditionally regarded as challenges that are solved mainly through cerebral activity, specifically via information-processing steps carried by neurons in the cerebral cortex. Activity in cortical networks thus constitutes a key factor for improving our understanding cognitive processes. However, recent evidence has shown that evolutionary older players in the central nervous system, such as brainstem’s ascending modulatory systems, might play an equally important role in diverse cognitive mechanisms. Our article examines the role of the ascending arousal system on large-scale network dynamics by combining pupillometry, functional MRI and graph theoretical analysis*.

## Introduction

Cognitive processes emerge from the dynamic interplay between diverse mesoscopic brain systems ^1,2^. Thus, the neural activity supporting cognition does not exist in a vacuum, but instead is deeply embedded within the ongoing dynamics of the physiological networks of the body^3^. In particular, the neural processes underlying cognition are shaped and constrained by the ascending arousal system, whose activity acts to facilitate the integration between internal states and external contingencies^4^. Timely and selective interactions between the ascending arousal system and the network-level configuration of the brain are thus likely to represent crucial constraints on cognitive and attentional processes. Yet, despite these links, we currently have a relatively poor understanding of how the ascending arousal system helps the brain as a whole to functionally reconfigure during cognitive processes, such as attention, in order to facilitate effective cognitive performance.

Recent evidence has linked higher-order cognitive functions in the brain to the intersection between whole-brain functional network architecture and the autonomic arousal system^2,5–8^. Central to these relationships is the unique neuroanatomy of the ascending noradrenergic system. For instance, the pontine locus coeruleus, which is a major hub of the ascending arousal system, sends widespread projections to the rest of the brain^9^. Upon contact, adrenergic axons release noradrenaline, which acts as a ligand on three types of post- and pre-synaptic adrenergic receptors (i.e., *α*1, *α*2 and β). The functional effects of each of these receptors depend on their differential sensitivities to noradrenaline (affinities for the ligand differ across receptors: *α*2 > *α*1 > β) and intracellular cascades, as well as their neuronal and regional distributions^9–14^. By modulating the excitability of targeted regions, the locus coeruleus can effectively coordinate neural dynamics across large portions of the cerebral cortex^15,16^. However, it is challenging to non-invasively track the engagement of the locus coeruleus during whole-brain neuroimaging and cognitive task performance.

Fortunately, it has been widely shown that the pupil diameter directly responds to changes in the activity of the locus coeruleus, and thus serves as an indirect, non-invasive measure of the noradrenergic system^17,18^.Specifically pupil diameter has been shown to indirectly monitor the neuromodulatory influences of the ascending arousal system on a variety of different brain regions^5,11,19–21^. Moreover, noradrenergic-mediated dilations in pupil diameter have been shown to effectively track the allocation of attentional resources^22–24^, in addition to both physical and mentally effortful processes^25,26^. Fast, phasic changes in pupil diameter have also been shown to directly relate to changes in the activity of the locus coeruleus^18,27,28^. While there is some evidence that pupil diameter covaries with other subcortical systems, such as the cholinergic^29^ and serotoninergic system^30^, the physiological mechanism for these effects is more opaque, and there is also clear causal evidence linking stimulation of the locus coeruleus to dilation of the pupil^19,31^. Despite these insights, several questions remain unanswered regarding how these processes are related to the complex architecture of the brain^32^. For instance, the processes by which the ascending arousal system modulates the functional dynamics of brain networks to facilitate attention, decision making and optimal behavioural performance have only begun to be explored^31,33–35^.

To examine these relationships in more detail, participants performed a motion-tracking task (top panel of Figure 1A) involving four levels of increasing attentional load, which was modulated by manipulating the number of items required to covertly attend to over an 11s tracking period. Specifically, subjects were instructed to covertly track the movement of several pre-identified targets (two to five) in a field of non-target stimuli (ten in total, including targets; see Figure 1). To investigate the network topological signatures of performing this task, we collected concurrent BOLD fMRI and pupillometry data. We hypothesized that, if increasing mental effort led to the reconfiguration of large-scale network architecture via the ascending arousal system, then the number of items required to be tracked over time (i.e., the attentional load) should relate to: i) increased pupil diameter; ii) heightened BOLD activity within attentional networks; and iii) augmented topological integration. Also, we predicted that individual differences in pupil diameter should track individual differences in effective attentional performance and decision processes^35–37^. Finally, we tested if the regional patterns of network configuration were predicted by the distribution of a predefined adrenergic receptor density atlas^31,34,38,39^. Our results confirm these predictions, and hence provide a mechanistic link between network topology, ascending noradrenergic arousal and attentional load.

**Figure 1:**
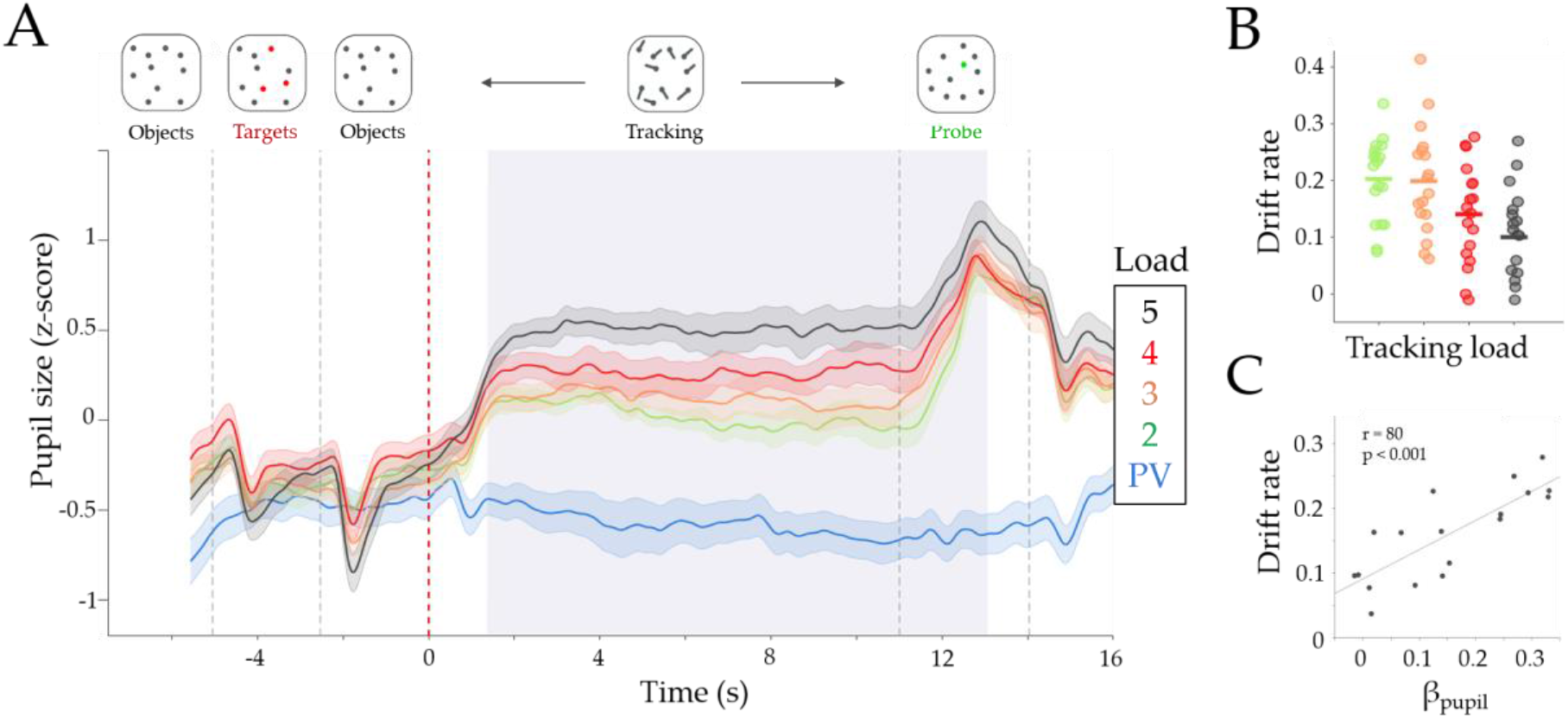
Effect of task difficulty on pupil diameter. A) Group average (z-score) pupil diameter time series for each Load condition. Colors represent passive viewing (PV) in blue, and Loads 2 to 5 in green, orange, red and black, respectively. The shaded area represents the standard error of the mean. We observed an average increase in pupil diameter, during tracking, with each Load condition. The light grey area represents timepoints with significant parametric effect (β_pupil_ > 0; FDR corrected at *p* < 0.01). Dotted lines represent the onset of each trial event (showed in the top part of the Figure). The red dotted line (Time = 0) is the tracking onset period when the dots began to move; B) Drift rate in each load condition. Each dot is the drift rate for each subject and load (mean β_Drift_ = −0.03, t_(17)_ = −7.43, *p* = 9.7×10^-7^); C) Pearson correlation between the pupil parametric effect of Load (β_pupil_) with the average drift rate across subjects (r_drift_ = 0.8, *p* = 1.0×10^-4^). The x-axis is the mean beta estimate of the pupillary load effect of the significative time window (β_pupil_) and the y-axis represents the mean drift rate across Loads.

## Results

### The Relationship Between Sympathetic Tone and Attentional Processing

Consistent with previous work^5^, our two level analysis - linear regression within each subject, and a two-tailed *t*-test between subjects - found that task performance (i.e., correct responses) decreased with attentional load (mean β_Acc_ = −6.66; t_(17)_ = −5.19, *p* = 7.2×10^-5^; Figure S1B) while RT increased with attentional load (mean β_RT_ = 0.06, t_(17)_ = 5.10, *p* = 8.8×10^-5^). We expanded on this result by translating performance into EZ-diffusion model parameters. Roughly, this approach uses the accuracy and reaction time distribution to estimate three latent parameters^40^: drift rate, a marker of the accumulation of decision evidence (Eq. 1); boundary criteria, the amount of evidence required to make a decision (Eq. 2); and non-decision time, the epoch spent processing the tasks perceptually (Eq. 3). The advantages of using this model are twofold: firstly, there are well-known links between the parameters to decision making processes^41,42^, pupil diameter^27,43^ and network reconfiguration^2^; secondly, drift rate accounts for the accuracy-reaction time trade off, as it takes into consideration both accuracy and the variability in reaction time into its calculation. In this way, our approach offers a better approximation of the ongoing computational processing during the task than accuracy and RT^44,45^. Using this approach, we observed a decrease in both the boundary criteria (β_Bound_ = −0.01, t_(17)_ = −2.70, *p* = 0.015) and drift rate (mean β_Drift_ = −0.03, t_(17)_ = −7.43, *p* = 9.7×10^-7^; Figure 1B), and an increase in the non decision time (mean β_nd_ = 0.07, t_(17)_ = 5.32, *p* = 5.5×10^-5^) with increasing attentional load.

By calculating the linear effect of load on pupil size across a moving average window of 160ms (see Methods), we observed a main effect of increased pupil diameter across both the tracking and probe epochs (β_pupil_ > 0, *p*_FDR_ < 0.01; light grey in Figure 1A depict significant epochs of time during the task; and in Figure S1A show the group average β_pupil_ time series). We also observed a positive correlation between mean β_pupil_ during the significant period (for simplicity we will refer to this value as βpupil) to the mean drift rate, mean boundary criteria and accuracy across all loads (Pearson’s r_drift_ = 0.8, *p* = 1.0×10^-4^; Figure 1C; r_acc_ = 0.68, *p* = 1.5×10^-3^, Figure S1C; r_*α*_ = 0.71, *p* = 9×10^-4^). The same relationships were not observed with non-decision time (Pearson’s r_nd_ = −0.31, *p* = 0.19). Additionally, we analysed whether this effect was present both within and between subjects in a trial-by-trial manner. To this end, we created a logistic linear mixed model (Eq. 6) to test whether pupil diameter was a predictor of performance (i.e., correct or incorrect response), as we would expect that incorrect responses should relate to decreased pupil diameter in difficult trials. We used the average pupil diameter within each trial of Load 4 and 5 (to account for the ceiling effect of Load 2 and 3) as regressors and subject as a grouping variable. We found a statistically significant fixed effect of pupil diameter on performance within each trial (β = 0.0127 ± 5×10^-4^; t_(286)_= 2.48; *p* = 0.013). Furthermore, we analyzed the random effect coefficients, which are the dispersion of the regressor across the grouping variable from the fixed regressor (in this case there is one value per subject), to assess the role of average across task performance. We found that the random effect covaried with the average performance and drift rate of each subject (Accuracy: Pearson’s r = 0.73, *p* = 8×10^-5^; Drift: Pearson’s r = 0.73, *p* = 5×10^-5^) suggesting that trial by trial pupil diameter was a better predictor of performance (i.e., correct or incorrect) on subjects with higher average performance in comparison to subjects with lower performance across the task. In conclusion, these results suggest that attentional load manipulation and pupil dilation covaried with performance on this attentionally demanding task both within and between subjects.

### Network Integration Increases as a Function of Attentional Load

Based on previous studies, we hypothesized that an increase in attentional load should recruit a distributed functional network architecture^5^, heightening network integration^2,12,34^. To test this hypothesis, we implemented a hierarchical topological network analysis^46–48^ on the average time-resolved functional connectivity matrix calculated across the tracking period of the task. Our analysis identified a subnetwork of tightly inter-connected regions that were part of attentional, somatomotor, and cerebellar network (red in Figure 2) that increased its BOLD activity after the tracking onset (Figure 2F). The tightly integrated regions were diversely connected to a separate frontoparietal sub-module (blue in Figure 2) that was less active during the trial. Two remaining sub-modules (yellow and green in Figure 2) showed a negative BOLD response during the tracking period and were part of a diverse set of networks. Interestingly, 81% of the Frontoparietal network (FPN) and all the Default Mode Network (DMN) were found to be within this less active group (see Supplementary Table S2 for the complete list of regions and sub-module assignments).

**Figure 2:**
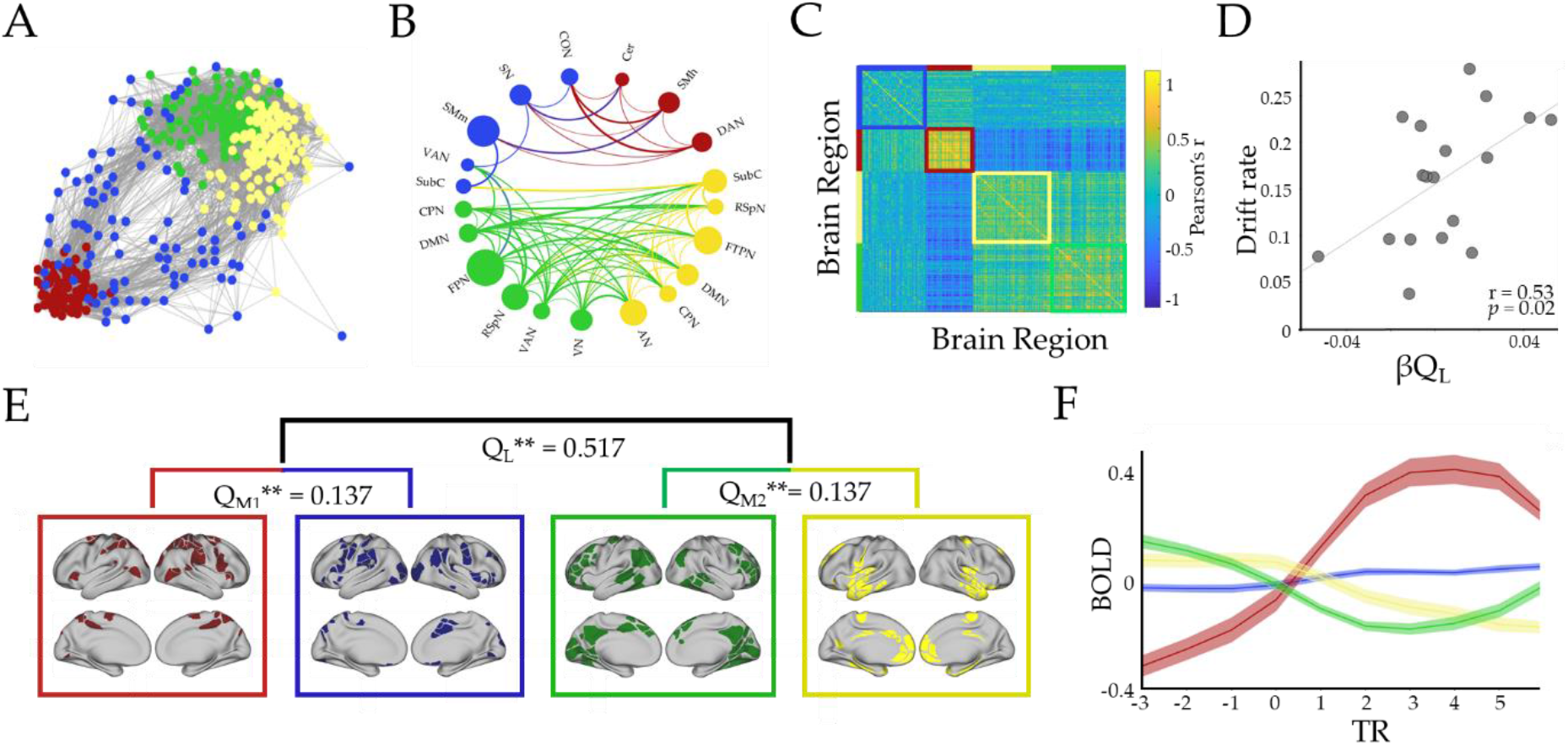
Hierarchical functional topology analysis of the brain during tracking across all loads. We observed two large-scale modules, and two meso-scale modules within each larger module (Module one [M1, red/blue] and Module two [M2, green/yellow], respectively): M1 corresponded to predominantly attentional and somatomotor network, and M2 to Frontoparietal (FPN) and Default Mode Network (DMN) among others (B and E). A) Forced directed plot representation of the average cluster across subjects. Edges higher than 0.15 are shown. Each color represents a unique sub module; B) A circle plot representing the resting state regions that were included within each sub module, with networks with > 30% of regions in each submodule shown in the plot. The diameter of the circles corresponds to the percentage of network regions that participated in that cluster. Connection width relates to average positive connection strength (functional connectivity), however only connections with r > 0.1 are shown; C) Connectivity matrix (Pearson’s r) between all pair of regions ordered by module assignments – note the strong anti-correlation between the red and green/yellow sub-modules; D) Correlation between parametric load effect on large scale modularity (β_Q_ value), and drift rate (Pearson’s r = 0.53; *p* = 0.022); E) Hierarchical analysis representation: *Q*_L_, *Q*_M1_ and *Q*_M2_ represent the modularity value for each level (*Q*_L_ large scale, and *Q*_M1-M2_ meso-scale level) and ** represents the probability of finding this value when running a null model (*p* = 0 for all three modularity values). The brain maps correspond to the cortical regions associated with each sub module; F) BOLD mean effect for each sub-cluster, each line represents the group average, and shaded areas are the standard error of the mean, x-axis is Repetition Time (TR) centered around tracking onset (TR = 0). DAN, dorsal attention; VN, visual; FPN, frontoparietal; SN, salience; CO, cingulo-opercular; VAN, ventral attention; SMm, somatomotor mouth; SMh, somatomotor hand; RSpN, retrosplenial; FTP, frontotemporal; DMN, default mode; AN, auditory; CPN, cinguloparietal; SubC, subcortex; Cer, Cerebellar.

Contrary to expectations, we did not observe significant parametric topological change (i.e., modularity, Q) at the macroscopic level as a function of attentional load (p > 0.05 for all TRs, Figure S2A). However, when analysing the correlation between modularity and performance measures (i.e., accuracy, drift rate and pupil diameter), we observed that an increase in the large-scale modularity load effect (i.e., higher modularity with load, β_QL_) positively correlated with higher mean drift rate (Pearson’s r = 0.53; *p* = 0.022; Figure 2D), mean accuracy (Pearson’s r = 0.61; *p* = 0.007; Supplementary Figure S3A), but was independent from βpupil (Pearson’s r = 0.43; *p* = 0.073). These results suggested that the system reconfigured during tracking towards increasing modularity, which in turn affected the efficient encoding of the ongoing task during tracking and hence, the decision-making process during the task probe.

Upon closer inspection of the data (Figure 2C), we observed a substantial number of nodes that were playing an integrative role during task performance, albeit at a finer resolution than the initial analysis suggested. We performed the modularity assignment within each large-scale module. The hierarchical analysis resulted in two pairs of sub-modules at the meso-scale level with a significant modularity (compared to 100 random graphs with preserved signed degree distribution; Q_M1_ = 0.137, *p* = 0; Q_M2_ = 0.137, *p* = 0; Figure 2E). Specifically, the red sub-module was found to selectively increase its participation coefficient (PC) at the meso-scale level (i.e., by increasing the connection weights to the blue sub-module in comparison to intramodular connections; Eq. 5) as a function of increasing attentional load (β_PC_ = 2.4×10^-3^, t_(17)_ = 3.57; *p* = 0.002; Figure 3A). Additionally, the extent of integration in the red sub-module was positively correlated across subjects with βpupil(Pearson r = 0.62, *p* = 0.006; Figure 3B), drift rate (Pearson’s r = 0.66, *p* = 0.002; Figure 3C) and accuracy (r = 0.57, *p* = 0.012, Figure S3B). Importantly, these relationships were found to be specific to the red submodule (Blue: Pearson’s r = −0.02, *p* = 0.936; Yellow: Pearson’s r = −0.011, *p* = 0.965; Green: Pearson’s r = −0.12, *p* = 0.617).

**Figure 3:**
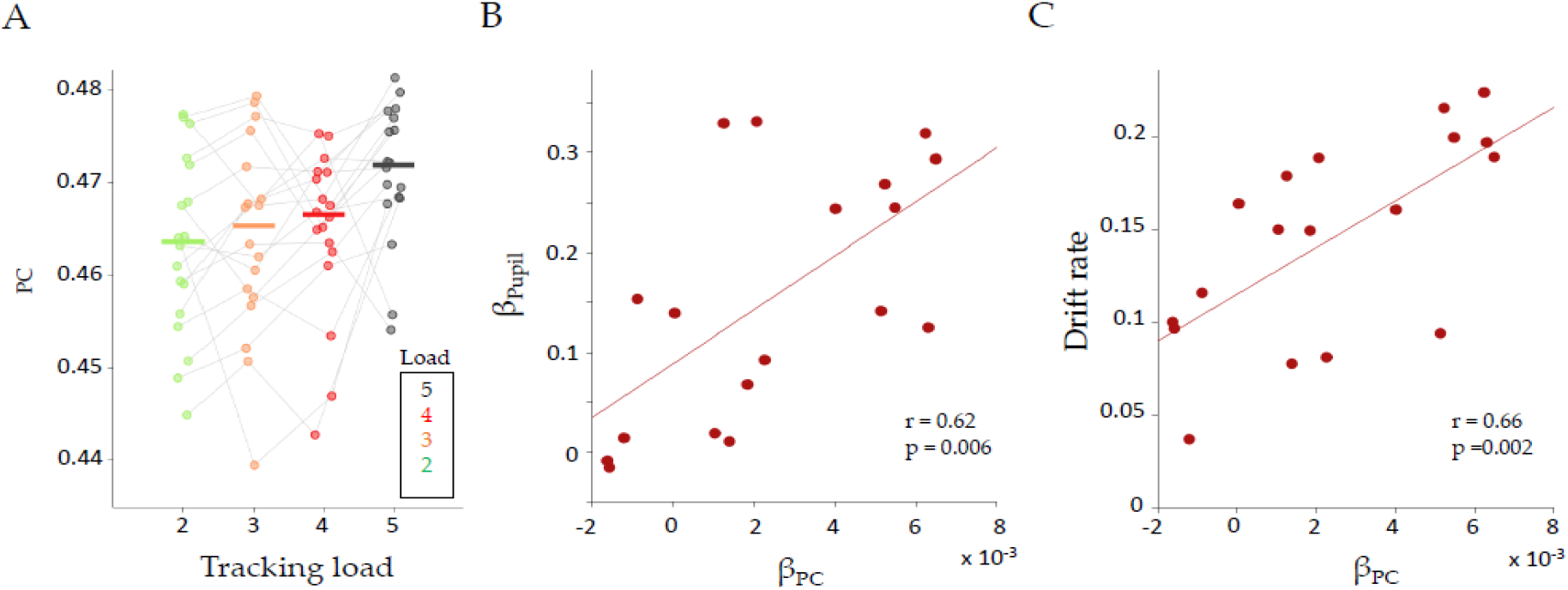
Relationships between load effect on participation, drift rate and pupil load effect. A) Average participation coefficient (PC) for each load, for the red module, during tracking. Each color represents the corresponding tracking load (from 2 to 5). Grey lines correspond to each subject; B-C) A regression parameter (β_PC_) was calculated for each subject and then correlated to β_pupil_ (B; r = 0.62; *p* = 0.006) and Drift rate (C; r = 0.66; *p* = 2.4×10^-3^). Each circle corresponds to the mean value per subject.

Based on these results, we implemented a linear mixed model (Eq. 7), using the subjects’ average pupil response within each Load as a regressor and the average participation of the red sub module as the dependent variable, with grouping by subject. Using this approach, we observed a significant fixed effect of pupil diameter on PC (β = 7.6×10^-3^ ± 3×10^-3^, *t*_(70)_ = 2.60, *p* = 0.011). Furthermore, the random effect coefficients (i.e., the between subject variation of the regressor value) correlated positively with accuracy (Pearson’s r = 0.47, *p* = 0.048) and drift rate (Pearson’s r = 0.62, *p* = 0.005), suggesting that subjects with a strong relationship between red module integration and pupil diameter have better behavioural outcomes. We then correlated the red βPC to the load effect on large scale modularity (βQ_L_, Fig. 2D) and observed a significant positive correlation (Pearson’s r = 0.59, *p* = 0.009). Finally, given that both of the topological parameters were correlated to drift rate and also with each other, we performed a partial correlation between drift rate and βPC controlling by βQ_L_ (r = 0.51, *p* = 0.034), and the partial correlation between drift rate and βQ_L_ controlling by βPC (r = 0.36, *p* = 0.145). This suggests that drift rate is correlated to the mesoscale integration of the red sub-module, but less so with increases in large scale modularity. Thus, although the macroscale network did not demonstrate increased integration *per se*, the relative amount of meso-scale integration within the red community was associated with increased performance (i.e., drift rate) and sympathetic arousal (i.e., pupil diameter), both between and within subjects. In this way, these results provide a direct relationship between the effect of attention load on pupillometry, drift rate, and a trade-off between large-scale segregation and meso-scale network integration.

### Network meso-scale integration and adrenergic receptor density

Given the relationship between mental effort, noradrenergic tone and pupil dilation^5,18,26,49,50^, the results of our analyses strongly suggested that the adrenergic system is involved in the meso-scale network reconfiguration observed during attentional tracking. The locus coeruleus can impact the cortical system in multiple ways, both through direct release of noradrenaline onto cortical neurons, and through the modulation of subcortical regions (such as the thalamic nuclei) with concurrent impact on the cortical dynamic. Importantly, in either case, the modulation is dependent on the noradrenergic receptors subtypes, which have different sensitivities to noradrenaline^13,51^, variable expression in the cerebral cortex^52,5,37,58^ and also belong to distinct classes (i.e., *α*1, *α*2, and *β* receptors). In particular, the *α*2a has been previously associated with working memory, adaptive gain and effective attention^13,51,54^. To gain a deeper insight into the role of *α*2a receptors in mesoscale integration during attentional tracking, we extracted the regional expression of the ADRA2A gene (which codes for *α*2a adrenoceptors) from the Allen Human Brain Atlas repository^55,56^, and compared the cortical regional expression of this gene with the brain activity patterns identified in our network analysis (Figure 2E).

Based on the relationships between pupil diameter (Figure 1), topological signatures (Figure 2) and task performance (Figure 3), and the known link between these variables and engagement of the noradrenergic system, we hypothesized that the different modules and sub-modules that we observed should have different densities of neuromodulatory receptors to account for the differential patterns across the network. To test this hypothesis, we conducted a two-tailed *t*-test in each hierarchical level comparing the density of the ADRA2A expression between modules. To account for spatial autocorrelation, we generated 5,000 surrogates maps with the same spatial autocorrelation of the ADRA2A map, calculated a *t*-statistic for each surrogate and the evaluated the probability of finding the observed *t*-statistic against the null distribution^57,58^. We indeed observed significant differences between modules at the meso-scale level. Specifically, we found significant differences between the blue and yellow sub-modules (*t*_(194)_ = 3.82, *p* = 2×10^-4^, *p*_SA_ = 0.02) and the differences between green and yellow sub-modules (*t*_(177)_ = −4.47, *p* = 1.3×10^-5^, *p*_SA_ = 0.004), while the other differences did not survive the spatial autocorrelation test (green-red: *t*_(152)_ = 0.47; *p* = 0.635, *p*_SA_ = 0.590; yellow-red: *t*_(156)_ = −3.02, *p* = 0.003, *p*_SA_ = 0.121; green-blue: *t*_(173)_ = −0.68, *p* = 0.496, *p*_SA_ = 0.324; red-blue: *t*_(135)_ = −1.30, *p* = 0.195, *p*_SA_ = 0.237; Figure S5A).

The modulatory effects of noradrenaline have been argued to depend directly on ongoing glutamatergic activity in target regions^59,60^. Moreover, it has been shown that the main source of the BOLD activity is the neurovascular response caused by pyramidal neurons containing Cyclo-oxygenase-2^61^. Importantly, this evoked response following noradrenergic activation is dependent on the ongoing activity of the pyramidal neurons^62^. Thus, the role of noradrenaline on brain dynamics and BOLD response depends critically on ongoing glutamatergic activity, which putatively represents pooled neural spiking activity^63^. Given the differential task-related BOLD activity of the different sub-modules (i.e., Figure 2F, Figure S4and Figure 4A), and the observed regional variability and specificity of integration across the network, we hypothesized that network-level integration would be explained by the combined effect of ongoing BOLD activity and the distribution of the adrenergic receptor expression. Finally, we predicted that the role of the *α*2a receptor atlas in shaping brain activity and topology should be dependent of the subjects’ pupil diameter, such that higher βpupil should rely on a stronger relationship between network topology and *α*2a receptor expression.

**Figure 4:**
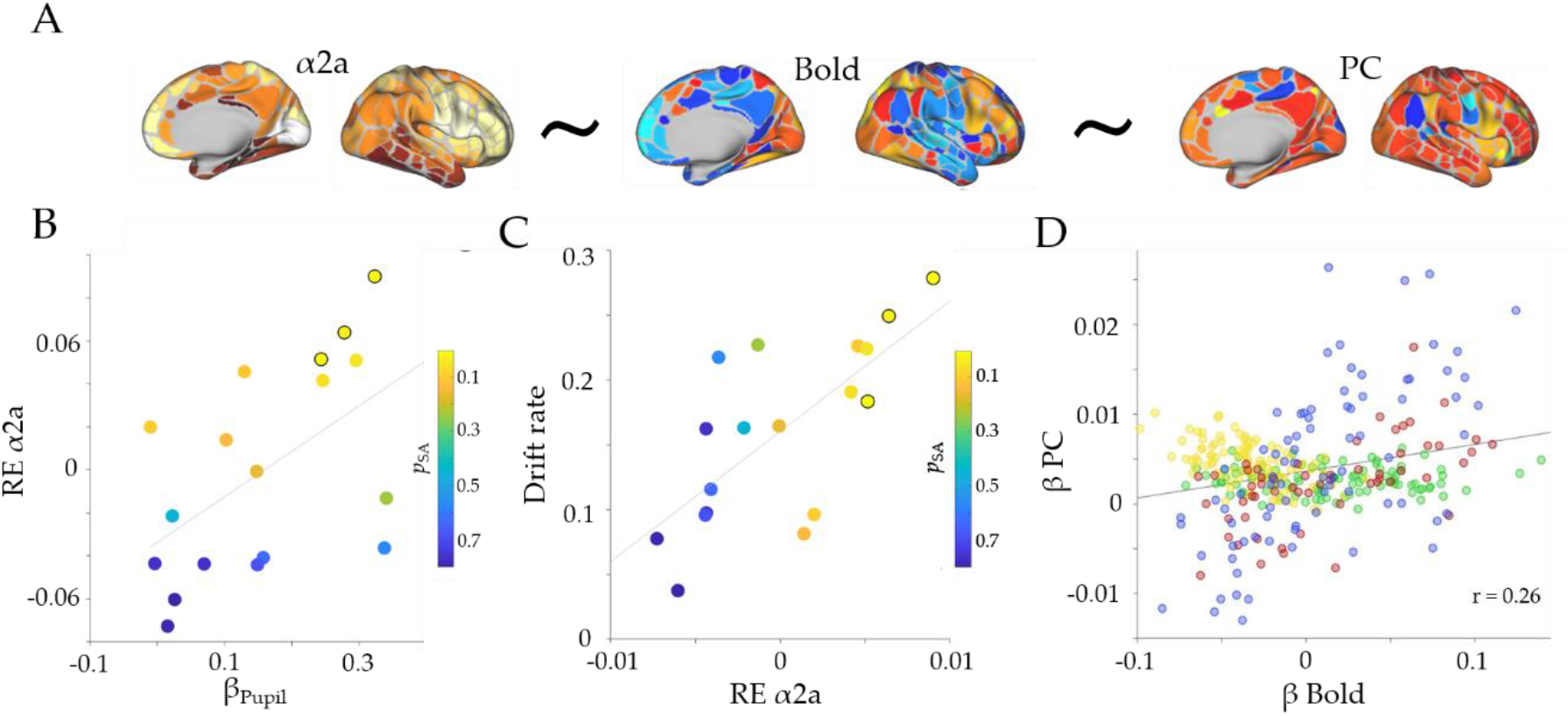
Receptor density analysis. A) Spatial maps of *α*2a density (left), BOLD parametric effect (middle) and Participation Coefficient parametric effect (right); ‘~’ represents the linear model tested in the analysis; B) Scatter plot depicting the relationship between β_pupil_ and the random effect of *α*2a (RE *α*2a; r = 0.54, *p* = 0.02); C) Scatter plot depicting the relationship between the random effect of *α*2a and drift rate (r = 0.70, *p* = 0.001) – the colors of the dots represent the *p*_SA_ value from the linear effect of *α*2a on βBOLD within each subject and the marked circles correspond to subjects with *p*_SA_ < 0.05; D) Pearson correlation of the group average BOLD parametric effect (β_BOLD_) and participation coefficient (β_PC_; r = 0.26, *p* = 17 x 10^-7^). Colors represent each module assignment as in Figure 2.

To evaluate between these different hypotheses, we created three linear mixed models in order to better disentangle the different plausible interactions between the variables (see Methods), while still controlling for between subject’s variability as grouping variable. Additionally, to control for spatial autocorrelation, we used 5,000 surrogate maps that maintained the spatial autocorrelation of the *α*2a while permuting the density values. In the first model (Eq. 8), we tested the hypothesis that the parametric BOLD effect (i.e., βBOLD, Figure S4) is shaped by the distribution of *α*2a receptors. We found significant evidence for a positive fixed effect of *α*2a on βBOLD activity, however this effect did not survive correction for spatial autocorrelation (β_*α*2a_ = 0.037± 0.016; t_(5992)_= 2.29; *p* = 0.022; *p_SA_* = 0.106; Table S2). Furthermore, we correlated the random effect coefficients (from the original and the surrogate maps) to both βPC and βpupil, and observed a significant positive correlation between the participation coefficient and both pupils (Pearson’s r = 0.54, *p* = 0.02, *p_SA_* = 0.036; Figure 4B) and mean drift rate (Pearson’s r = 0.70, *p* = 0.001, *p_SA_* = 0.001; Figure 4C). This result shows the manner in which pupil diameter linearly shapes βBOLD cortical map through the engagement of the *α*2a receptor expression map. Importantly, although the fixed effect of *α*2a on βBOLD didn’t survive the spatial autocorrelation correction, the linear correlation of this effect with both βpupil and drift rate (between subjects) did survive the correction.

To further analyze the between subject differences in the role of *α*2a receptor atlas in shaping the βBOLD map, we ran a separate linear model within each subject with *α*2a as a regressor and βBOLD of each region as the dependent variable (while also correcting for spatial autocorrelation using 5,000 surrogate maps). As can be seen in Figure 4B-C, we observed a dependency between the *p_SA_* value, βpupil and drift rate, in which the βpupil and drift rate subject effects survived the spatial autocorrelation correction (*p_SA_* < 0.05; marked circles in Fig. 4B-C). Despite these results, there was no significant effect of *α*2a on βPC (Eq. 9; β_*α*2a_ = 0.001 ± 0.003; t_(5992)_= −0.51; *p* = 0.6), and no significant Pearson’s correlation were found between the random effects and both β_pupil_ or drift rate (r = −0.24, *p* = 0.33 and r = −0.23, *p* = 0.341, respectively). However, we did find a significant effect of βBOLD on βPC (Eq. 10; β = 0.0259 ± 0.006; t_(5992)_ = 3.96; *p* = 7.55 x 10^-5^). Together these results propose a closer link between pupil diameter, ascending neuromodulation and the cortical neuromodulation dependent on *α*2a receptor density.

Finally, we observed a differential relationship between βPC and βBOLD depending on the large-scale module to which the regions were assigned (Figure 4D). We expanded the former result by measuring, within each subject, the Pearson correlation between the β_BOLD_ and β_PC_ separately in each module (M1 being the modules assigned as red and blue, and M2 assigned as yellow and green; Figure 2). The results demonstrated a significant difference between modules, meaning that the M1has a higher correlation with βPC, in comparison to M2 (t_(17)_ = −12.99, *p* = 2.93 x 10^-10^). These results provided evidence that the adrenergic receptor distribution of *α*2a shapes the βBOLD activation map in proportion to the subject’s pupil diameter. Additionally, βBOLD activation map modulates (i.e., was related to) meso-scale integration, and meso-scale integration is related to pupil diameter. Based on these results, we hypothesise that the adrenergic system shapes the BOLD activity, which in turns shapes the topology of the network towards integration. However, future work is required in order to test this hypothesis more directly, for instance by combining optogenetic approaches with neuronal recordings in awake animals.

## Discussion

Here, we leveraged a unique dataset to simultaneously track pupil diameter and network topology during an attentional demanding task with increasing attentional load. Our results provide integrative evidence that links the ascending arousal system to the mesoscale topological signature of the functional brain network during the processing of an attentionally demanding cognitive task. Pupil diameter tracked with attentional load (Figure 1A) and was related to the speed of information accumulation as estimated by a drift diffusion model (Figure 1B-C). Additionally, we observed concurrent pupil dilations and adaptive mesoscale parametric topological changes as a function of task demands (Figures 2 and 3). Finally, we found evidence that topological reconfiguration was dependent on the regional activity and the genetic expression of the adrenergic receptors in the brain (Figure 4). Together, these results provide evidence for the manner in which the ascending arousal noradrenergic system reconfigures brain network topology so as to promote attentional performance according to task demands.

The relationship between performance and pupil diameter is consistent with the predictions of Adaptive Gain Theory^17^. Within this framework, the locus coeruleus is proposed to adaptively alter its activity according to the demands imposed on the system. More specifically, the theory proposes that performance follows an inverted U-shaped relationship with arousal, such that maximal operational flexibility in the noradrenergic system is associated with optimal task performance^13,54^. We observed that load-related increases in pupil diameter, presumably due to increased activity in the ascending arousal system^17,18,64^, relates closely with the activity and topology of the broader brain network (Figure 2), in a manner that is reflective of effective task performance (Figure 3). Similar effects have been described in animal models after a chemogenetic activation of the locus coeruleus, which strongly alters the large-scale network structure towards large-scale integration, specifically in regions with heightened adrenergic receptor expression^31^. How these changes, which are likely related to the modulation of the neural gain that mediates effective connections between distributed regions of the brain^15,33^, are traded-off against requirements for specificity and flexibility remains an important open question for future research.

The addition of attentional load was found to alter the integration of meso-scale sub-modules, but not the higher-level modular organization. This topological result is somewhat more targeted than those described in previous work^2,34,65^. While these differences may be related to disparities in the way that the data were analyzed, the results of our study do demonstrate that alterations in the cerebral network topology at a relatively local (i.e., sub-modular) level are crucial for effective task performance^66^. Additionally, our results replicate and expand upon a previous study^67^, in which the authors found that short term practice on an attentional task was related to increased coupling between attentional networks and segregation among task-negative (DMN) and frontoparietal network (FPN). Our study replicates the graph theoretical results of that study, while also directly relating the findings to the architecture of the ascending neuromodulatory system. One potential explanation for these results comes from animal studies, in which rapid changes in pupil diameter have been compared to changes in neural population activity at the microscale^18,49,50^. These studies suggest that the ascending arousal system may be able to alter the topology of the network in a hierarchical manner that is commensurate with the spatiotemporal scale of the arousal systems’ capacity ^2^. Future work that integrates results across spatiotemporal scales is required to appropriately adjudicate the implications of this hypothesis.

Importantly, our approach is not without limitations. For one, the participation measures used in our linear mixed model were estimated at the meso-scale level, and hence derived from different modular partitions. Furthermore, the specificity of the pupillary response as a correlate of LC activity is currently under active debate. For instance, in addition to the strong empirical links between the noradrenergic system and pupil dilation, there is also evidence that the pupil is dilated in concert with activity in the basal forebrain cholinergic system^68^, however it bears mention that both peripheral^69^ and central cholinergic tone^21^ is associated with pupillary constriction. There are more plausible physiological routes for the serotonergic system to dilate the pupil (via the excitation of the intermediolateral cell column), and in keeping with this, there is evidence that the serotonergic system is linked with pupil dilation^30^. Nevertheless, it is important to take into account that the neuromodulatory arousal system is replete with complex interconnections^70–73^. In addition, based on the current lack of a specific mechanism involving pupillary changes through the cholinergic system, it is highly probable that those correlations are due to indirect modulation of pupillary responses (e.g., via indirect neuromodulation mediated by the LC system). On another hand, we acknowledge the limitations of the atlas receptor analysis and the linear model used in our study. More specific neurobiological properties of the receptor distributions are needed to make better inferences, and hence provide more accurate answers of their role in brain dynamics. For instance, it would be ideal to compare receptor distributions that incorporated layer-specific expression, as there are well-known cellular and circuit differences across layers in the cerebral cortex^74,75^. Importantly, taking into consideration the strong correlation between different genetic expression maps^76^, it is possible that the current correlation between ADRA2A expression and brain activity is a false positive caused by another neuroanatomical gradient strongly correlated to the ADRA2A. Therefore, future work studying the interaction between genetic expression of the neuromodulatory receptors, pupil diameter and brain activity is needed. In spite of this limitation, we believe in the importance of integrating pupil diameter and receptor distribution in the analysis as the relationships between noradrenergic tone, brain activity and network topology will help us to disentangle the mechanistic steps connecting the locus coeruleus system to both pupil diameter and brain dynamics.

In summary, we provide evidence linking mesoscale topological network integration, hierarchical organization and BOLD dynamics in the human brain that increases in attentional load, thus providing further mechanistic clarity over the processes that underpin the Adaptive Gain Model of noradrenergic function in the central nervous system.

## Methods

### Participants

18 right-handed individuals (age 19–26 years; 5 male) were included in this study. Exclusion criteria included: standard contraindications for MRI; neurological disorders; mental disorders or drug abuse. All participants gave written informed consent before the experiment.

### Parametric Motion Tracking Task

Each trial of the task involved the same basic pattern (Figure 1A): the task begins with a display presenting the objects (i.e., blue colored disks); after a 2.5 s delay, a subset of the disks turns red for another 2.5 seconds; all of the disks then return to blue (2.5 seconds) before they started moving randomly inside the tracking area. The participants’ job is to track the ‘target’ dots on the screen while visually fixating at the cross located at the center of the screen. After a tracking period of ~11 seconds, one of the disks is highlighted in green (a ‘probe’) and the subject is then asked to respond, as quickly as possible, as to whether the green probe object was one of the original target objects. The number of objects that subjects were required to attend to across the tracking period varied across trials. There were five trial types: passive viewing (PV), in which no target is assigned; and four load conditions, in which two to five targets were assigned for tracking. We operationalized attentional load as the linear effect of increasing task difficulty (i.e., the number of targets to be tracked).

The experiment was conducted using a blocked design, in which each block included: instruction (1s); fixation (0.3s, present throughout the rest of trial); object presentation (all objects were blue; 2.5s); target assignment (i.e., the targets changed color from blue to red; 2.5s); object representation (objects back to the original blue color; 2.5s); object movement/attentional tracking (moving blue dots; 11s); object movement cessation (0.5s); and a final probe (color change to green and response; 2.5 s). The total duration of each trial was 22.8s. Each condition was repeated 4 times in one fMRI-run, which also included 4 separate fixation periods of 11s each between five consecutive trials. All participants completed 4 separate runs of the experiment, each of which comprised 267 volumes. The order of the conditions was pseudo-random, such that the different conditions were grouped in sub-runs of triplets: PV, pseudo-random blocks of Loads 2 through 5 and a fixation trial. All objects were identical during the tracking interval and standard object colors were isoluminant (to minimize incidental pupillary responses during the task).

### Behavior and EZ-Diffusion Model

The EZ-diffusion model was used to interpret the performance measures from the task^45,77^. This model considers the mean RT of correct trials, SD-RT across correct trials, and mean accuracy across the task and computes from these a value for drift rate (*v*, equation 1), boundary separation (*a*, equation 2), and non-decision time (equation 3) – the three main parameters for the drift-diffusion model^77,78^.

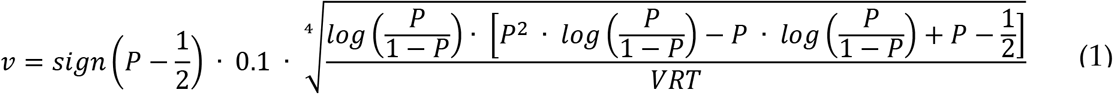

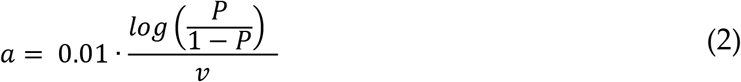

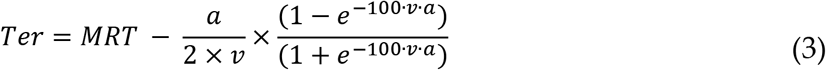

In which *P* is the average performance (range between 0 to 1); sign is an operator that will be −1 if *P* < 0.5 or +1 if *P* > 0.5; VRT is the standard deviation of reaction time (in seconds); and MRT is the mean reaction time (in seconds).

### Pupillometry

Fluctuations in pupil diameter of the left eye were collected using an MR-compatible coil-mounted infrared EyeTracking system (NNL EyeTracking camera, NordicNeuroLab, Bergen, Norway), at a sampling rate of 60 Hz and recorded using the iView X Software (SensoMotoric Instruments, SMI GmbH, Germany). Blinks, artifacts and outliers were removed and linearly interpolated^79^. High frequency noise was smoothed using a 2^nd^ order 2.5 Hz low-pass Butterworth filter. To obtain the pupil diameter average profile for each level of attentional load (Fig. 1B), data from each participant was normalized across each task block (corresponding to the five consecutive trials between fixations). This allowed us to correct for low frequency baseline changes without eliminating the load effect and baseline differences due to load manipulations^80,81^. Following this, a linear regression was performed in each time point using the task load as regressor and resulting in a ‘load effect’ time series for each subject.

### MRI Data

Imaging data were collected on a Philips Achieva 3 Tesla MR-scanner, equipped with an 8-channel Philips SENSE head coil (Philips Medical Systems, Best, Netherlands) at the Interventional Centre, Oslo University Hospital, Norway. Functional data were collected using a BOLD-sensitive T2* weighted echo-planar imaging sequence (36 slices, no gap; repetition time (TR), 2,2s; echo time (TE), 30 ms; flip-angle, 80°; voxel size, 3×3×3; field of view (FOV), 240×240 mm; interleaved acquisition). Anatomical T1-weighted images consisting of 180 sagittal oriented slices were obtained using a turbo field echo pulse sequence (TR, 6.7 ms; TE, 3.1 ms; flip angle 8°; voxel size 1×1.2×1.2 mm; FOV, 256×256 mm).

### fMRI Data Preprocessing

After realignment (using FSL’s MCFLIRT), we used FEAT to unwarp the EPI images in the y-direction with a 10% signal loss threshold and an effective echo spacing of 0.333. Following noise-cleaning with FIX (custom training set for scanner, threshold 20, included regression of estimated motion parameters), the un-warped EPI images were then smoothed at 6 mm FWHM, and non-linearly co-registered with the anatomical T1 to 2 mm isotropic MNI space. Temporal artifacts were identified in each dataset by calculating framewise displacement (FD) from the derivatives of the six rigid-body realignment parameters estimated during standard volume realignment^82^, as well as the root mean square change in BOLD signal from volume to volume (DVARS). Frames associated with FD > 0.25mm or DVARS > 2.5% were identified, however as no participants were identified with greater than 10% of the resting time points exceeding these values, no trials were excluded from further analysis. There were no differences in head motion parameters between the four sessions (p > 0.500). Following artifact detection, nuisance covariates associated with the 6 linear head movement parameters (and their temporal derivatives), DVARS, physiological regressors (created using the RETROICOR method) and anatomical masks from the CSF and deep cerebral WM were regressed from the data using the CompCor strategy^83^. Finally, in keeping with previous time-resolved connectivity experiments^84^, a temporal band pass filter (0.0071 < f < 0.125 Hz) was applied to the data.

### Brain Parcellation

Following pre-processing, the mean time series was extracted from 375 predefined regions-of-interest (ROI). To ensure whole-brain coverage, we extracted: 333 cortical parcels (161 and 162 regions from the left and right hemispheres, respectively) using the Gordon atlas^85^, 14 subcortical regions from Harvard-Oxford subcortical atlas (bilateral thalamus, caudate, putamen, ventral striatum, globus pallidus, amygdala and hippocampus; http://fsl.fmrib.ox.ac.uk/), and 28 cerebellar regions from the SUIT atlas^86^ for each participant in the study.

### Time-Resolved Functional Connectivity and Network Analysis

Following pre-processing, the mean time series was extracted from 375 predefined regions-of-interest (ROI). To estimate functional connectivity between the 375 ROIs, we used the Jack-knife correlation approach (JC)^87^. Briefly, this approach estimates the static correlations between each pair of regions, and then recalculates the correlation between each pair after systematically removing each temporal ‘slice’ of data (i.e., each TR). By subtracting the jack-knifed correlation matrix from the original ‘static’ matrix, the difference in connectivity at each slice from the static connectivity value can be used as an estimate of time-resolved functional connectivity between each pair of regions at each TR in a way that does not require windowing.

### Community Structure

The Louvain modularity algorithm from the Brain Connectivity Toolbox (BCT)^88^ was used in combination with the JC to estimate both time-averaged and time-resolved community structure. The Louvain algorithm iteratively maximizes the modularity statistic, *Q*, for different community assignments until the maximum possible score of *Q* has been obtained (equation 1).

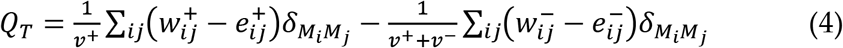

**Equation 1** – Louvain modularity algorithm, where *v* is the total weight of the network (sum of all negative and positive connections), *w_ij_* is the weighted and signed connection between regions *i* and *j*, *e_ij_* is the strength of a connection divided by the total weight of the network, and *δ_MiMj_* is set to 1 when regions are in the same community and 0 otherwise. ‘+’ and ‘−’ superscripts denote all positive and negative connections, respectively.

For each subject, we calculated the mean adjacency matrix from 1 TR before tracking until the end of the tracking period. Afterwards, a consensus partition was estimated across subjects. Finally, to identify multi-level structure in our data, we repeated the modularity analysis for each of the modules identified in the first step^46,47^. With this final module assignment, we were afforded an estimate of the time resolved, multi-level modularity (Q_*T*_) within each temporal window for each participant in the study.

### Regional Integration

Based on the group consensus community assignments, we estimated between-module connectivity using the participation coefficient, B_*T*_, which quantifies the extent to which a region connects across all modules (i.e. between-module strength; equation 2). In our experiment, we used two separate community assignments, one for each of the modularity levels. In this manner we measure: 1) how the first hierarchical level (i.e., large scale) topology changed during tracking across the complete brain; and 2) how the topology of the sub-modules changed across the task. These values were calculated in each time point using the time-resolved adjacency matrix across each load condition.

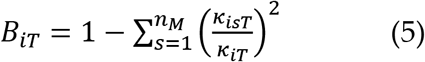

**Equation 2** - Participation coefficient B_*iT*_, where κ_isT_ is the strength of the positive connections of region *i* to regions in module *s* at time *T*, and κ_iT_ is the sum of strengths of all positive connections of region *i* at time *T*. The participation coefficient of a region is therefore close to 1 if its connections are uniformly distributed among all the modules and 0 if all of its links are within its own module.

### Neurotransmitter Receptor Mapping

To investigate the potential correlates of meso-scale integration, we interrogated the neurotransmitter receptor signature of each region of the brain. We used the Allen Brain Atlas micro-array atlas dataset (http://human.brain-map.org/)^55^ to identify the regional signature of genetic expression of the *α*2a subtype of the adrenergic receptor (ADRA2A). This receptor has been *a priori* related to cognitive function and attention^89^, and is one of the most abundant adrenergic subtypes expressed in the cerebral cortex^90^. This atlas contains postmortem samples of six donors that underwent microarray transcriptional characterization. The spatial map of *α*2a mRNA expression was obtained in volumetric 2mm isotropic MNI space, following improved nonlinear registration and whole-brain prediction using variogram modeling^91^. We used this data instead of the native sample-wise values in the AHBA database to prevent bias that could occur due to spatial inhomogeneity of the sampled locations. We projected the volumetric *α*2a expression data onto the Gordon atlas with linear interpolation and calculated the mean value within each parcel using custom MATLAB codes.

### Statistical analysis

#### The Relationship Between Sympathetic Tone and Attentional Processing

We analysed the between subjects’ effect of load on the behavioural, pupillometric and fMRI related variables by performing a two-level linear model analysis. In the first level, we used attentional load as a regressor (2 to 5) and -in independent models-the mean accuracy, reaction time, standard deviation of reaction time, drift rate, boundary criteria and non-decision time as dependent variables (i.e., 4 values per subject). From this, we ran a two-tailed t-test on the statistical effects (i.e., the β value from the regression, one for each subject; N = 18). Similarly, to calculate the load effect on pupil diameter, we calculated the average pupil diameter on each load condition within each subject. Then, we performed a first-level analysis in which we ran a linear regression in each time frame (1600 frames in total, corresponding to 26.6 seconds). This procedure resulted in one β timeseries (i.e., the statistical load effect on pupil diameter) for each subject across the trial (Figure S1A). After this, we performed a right tailed t-test in each frame across subjects (n = 18 in each frame) to find the periods of time where the β value where higher than zero. Finally, we corrected by false discovery rate (FDR)^92^ for multiple testing, which resulted in a period of time where the load effect was higher than 0 (light grey area in Figure 1A). The mean β values during this section was calculated in each subject and defined as ‘βpupil’. Finally, following the same pipeline, we calculated the effect of attentional load on the brain related signals (i.e., BOLD, participation coefficient [PC] and modularity [Q]). The effect of load on BOLD was calculated running a separate linear model in each subject and region within each TR (18 subjects; 375 regions; 10 TRs; 4 load condition), resulting in a matrix of β values of 18 x 375 x 10.

To evaluate the statistical effect of pupil diameter on accuracy, we performed a logistic linear mixed effects model. We used the mean pupil diameter of the significant time period (Figure 1A) of the high load trials (Load 4 and 5), and the accuracy (i.e., correct or incorrect) as the predictor variable of each trial, grouping by subject as the random effect. The statistical model is described in the following equation:

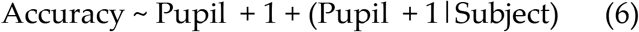

#### Network meso-scale integration and adrenergic receptor density

To evaluate whether the modularity of the network we observed was higher than chance, we generated 100 random networks in each hierarchical level (300 random networks in total), with a preserved degree distribution (using the MATLAB *randmio_und_signed* function from the Brain connectivity toolbox^88^). We calculated the modularity value of each random network and used the resultant values to populate a null distribution (Figure 2D).

We analyzed the statistical effect of pupil diameter on the participation coefficient both within and between subjects by performing a linear mixed model using the time varying PC of the red sub-module (Figure 3A) of each load as a dependent variable (N=72), and the respective pupil diameter as a regressor, with grouping by subject. The statistical model is described in the following equation:

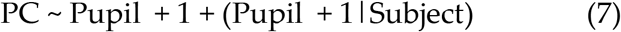

#### Network meso-scale integration and adrenergic receptor density

Expression of brain genetic atlas vary smoothly across the surface and thus is associated with non-trivial spatial autocorrelation that in turn violates the assumption of independence between samples^57,58,93^. To account for the spatial autocorrelation in these brain maps, we used spatial autocorrelation null maps as implemented in Brain Surrogate Maps with Autocorrelated Spatial Heterogeneity (BrainSMASH) python toolbox^57^. A geodesic distance matrix of the atlas parcels using the surface of the Gordon atlas was obtained to build the surrogates using BrainSMASH functions. We generated 5,000 null maps which were used to generate null distribution of the different statistics corrected by spatial autocorrelation.

We measure the statistical difference in the receptor density between sub-modules by a two-tailed t-test between each pair of modules. The same procedure was performed using the surrogate maps to generate a null distribution of t-statistics. To evaluate the effect of the density of each adrenergic receptor on the neural activity in the attentional task, we built a linear mixed model aimed at predicting regional differences in BOLD activity and participation coefficient. We created a model using the receptor density atlas of *α*2a receptor to predict parametric BOLD activity (i.e., linear increase of BOLD activity with task load) during tracking (Eq. 8). To evaluate the relationship between BOLD activity, adrenergic receptor expression and changes in participation coefficient as a function of attentional load, we tested two models: one using the adrenergic receptor density as independent factor (Eq. 9); and another using the parametric BOLD effect as an independent factor (Eq. 10). Additionally, we assessed the across-subject variability using the subjects ID as grouping variable in order to evaluate the random effects on the independent factor. We corrected the spatial autocorrelation by running the same model using 5,000 surrogate maps. Then we used the fixed effect null distribution to calculate the *p*_SA_ (i.e., the probability of finding the fixed effect within the 95^th^ percentile of the null distribution). The deterministic part of the model is expressed in the following equations^94^:

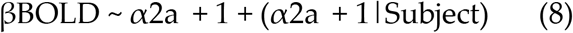

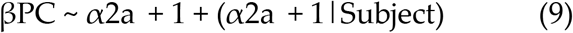

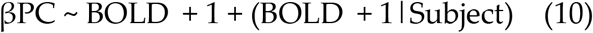

Where *PC* is the parametric effect of meso-scale participation coefficient (i.e., βPC), BOLD is the parametric effect of load on BOLD activity during tracking for each region, and *α*2a are the regional densities of the respective adrenergic receptor atlas. We then correlated the random effects parameters to pupil diameter responses and behaviour and then compared these with the Pearson’s correlation of the null distribution using the random effect of the surrogate maps. Finally, we performed a linear model within each subject with *α*2a as a regressor and βBOLD as dependent variable. Again, the statistical effect (i.e., β value) was compared against the null distribution when performing the regression using the surrogate maps (figure 4B-C).

## Data and code availability

The anonymized preprocessed fMRI and pupillometry data can be found at https://figshare.com/articles/dataset/MOT_data_mat/13244504. The ADRA2A expression atlas can be downloaded from http://www.meduniwien.ac.at/neuroimaging/mRNA.html. All analysis of the fMRI and pupil diameter data were performed on MATLAB 2020a. The surrogate maps of the ADRA2A atlas were generated on python. Documented code for reproducing the analyses is provided in https://github.com/gabwainstein/MOT.

## Acknowledgements, Funding and Disclosure

We thank P. Billeke for their thoughtful comments on our manuscript. JMS was supported by the University of Sydney Robinson Fellowship and NHMRC GNT1156536. GW was supported by ‘Becas Chile’ PhD scholarship. The authors declare no financial interests or conflicts of interest.

## Author Contributions

GW and JS Analysed the data, interpreted the results and wrote the manuscript. DR: Interpreted the results and wrote the manuscript. KK, DA, BL and TE: Created the experimental design and contributed with the data acquisition. All authors reviewed, commented and edited the manuscript, and all authors gave final approval of the version to be published.

