## Supplementary Information for "The ascending arousal system promotes optimal performance through meso-scale network integration in a visuospatial attentional task"

### Affiliations

### Corresponding author

### Supplementary Figures

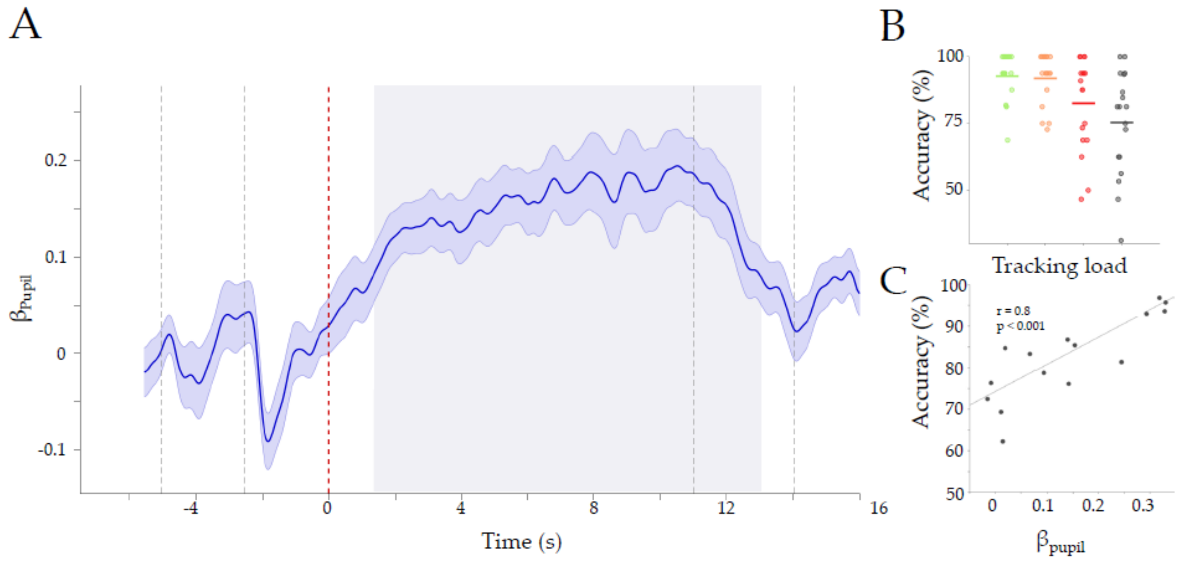


**Supplementary Figure S1:** A) regression parameter of the pupil diameter ($\beta_{Pupil}$) – grey area represents the significant time period where the group $\beta_{Pupil} > 0$ (FDR corrected, $p_{FDR} < 0.01$); B) Accuracy across loads, each dot is the group average. Colors represent Load 2 to 5 in green, orange, red and black, respectively; C) Pearson correlation between the average $\beta_{Pupil}$ during the significant period (grey area in A), to the average Accuracy. There is a significant positive correlation (Pearson r = 0.8, $p = 7.0\text{x}10^{-5}$).

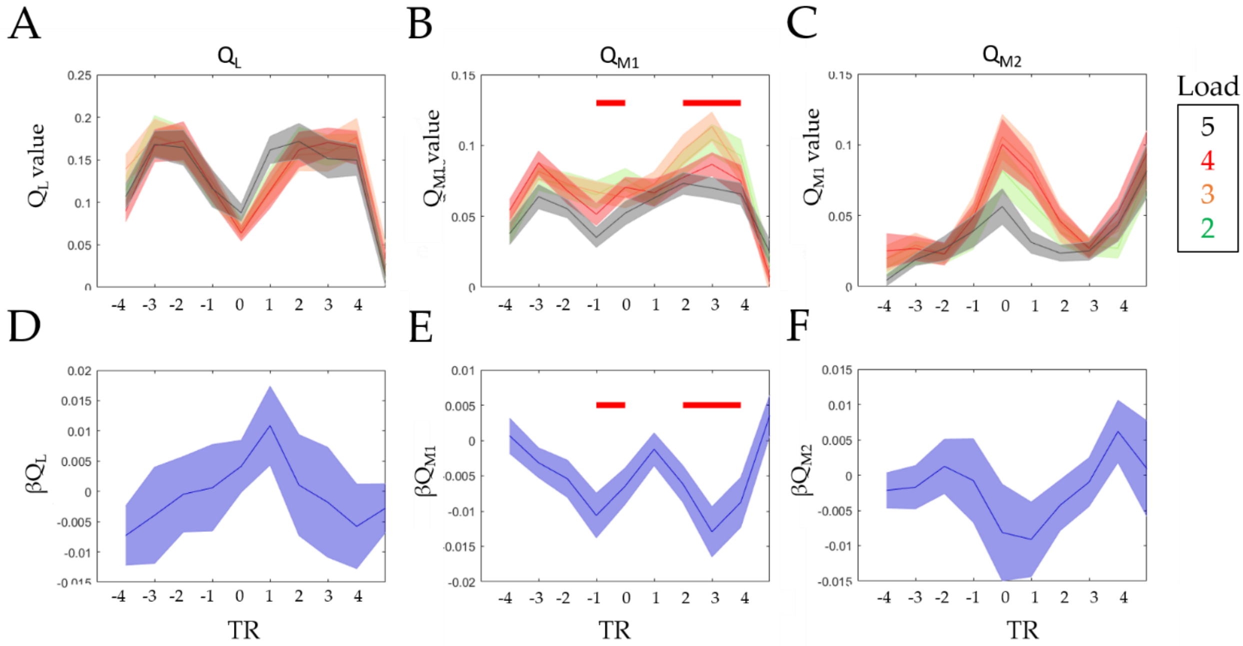


**Supplementary Figure S2:** Modularity time series for each attentional load and the parametric change across loads. In red (B and E) we show the significant time windows ($p_{FDR} < 0.05$, $\beta \neq 0$), which were only found for the $Q_{M1}$ sub module (module red/blue in the main text). x-axis is Repetition Time (TR) from tracking onset (TR = 0). Lines represent the group average, and the shaded area is the standard error of the mean. Colors in A-C represent Load 2 to 5 in green, orange, red and black, respectively.

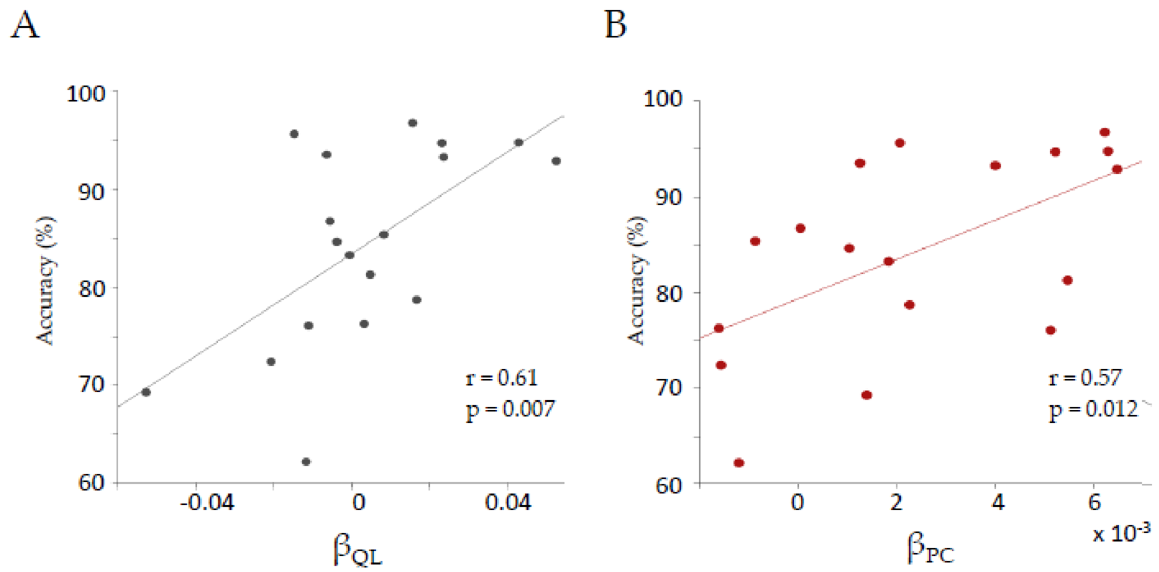


**Supplementary Figure S3:** Topological parametric effect ($\beta_{QL}$ and $\beta_{PC}$) correlation to Accuracy (A; Pearson r = 0.61, P = 0.007) and (B; Pearson r = 0.57, P = 0.012).

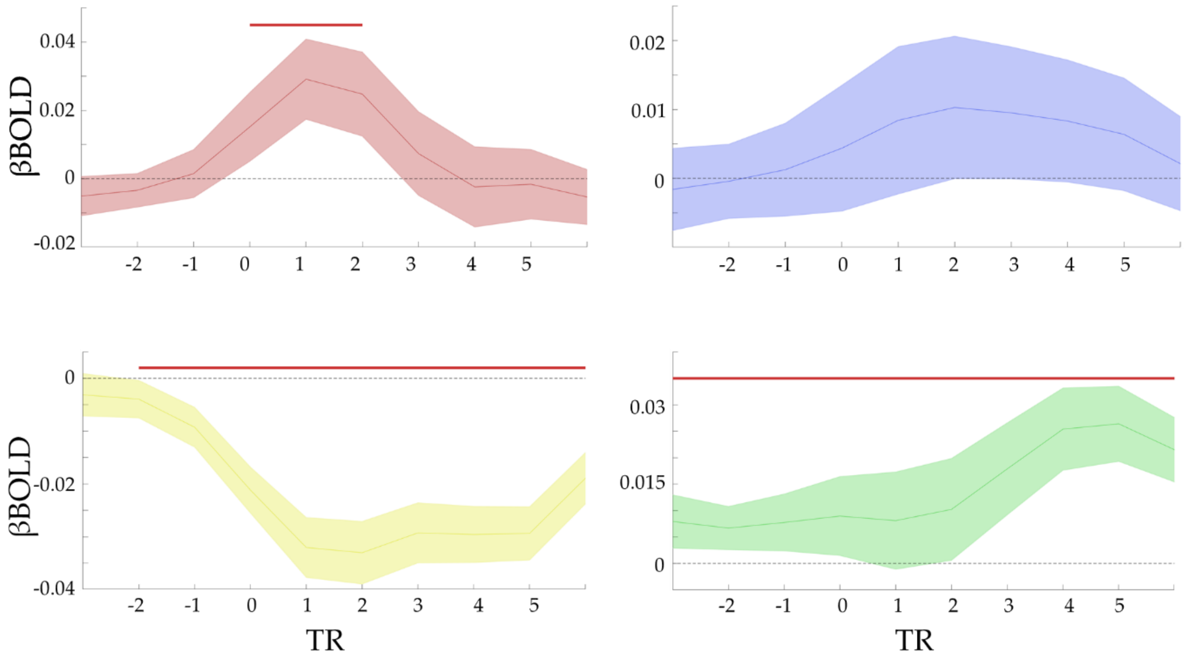


Supplementary Figure S4: Effect of attentional load on Bold activity. The statistical effect of load on bold is shown (βBOLD) for each TR (0 marks the start of tracking). Each plot represents the mean activity of each module. The colors represent the module assignment as in Figure 2 in main text. The red line in each plot shows the significant period of a two-tailed *t*-test ($\beta \neq 0$, $p_{FDR} < 0.05$) corrected by FDR.

.

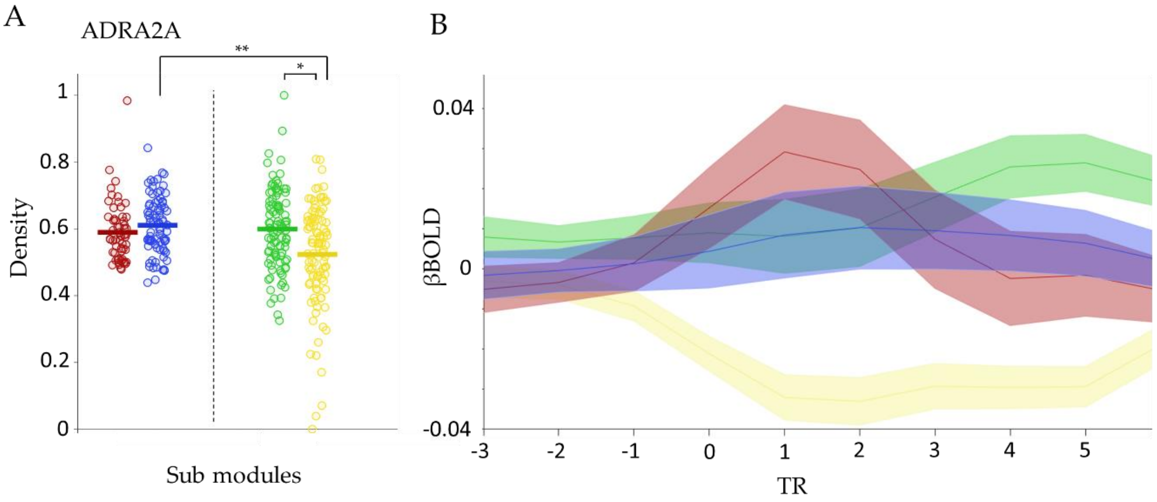


**Figure S5: A)** Receptor density of $\alpha$2a. The standardized density of the ADRA2A receptor expression, for each of the sub modules is shown. A two-tailed t-test was performed comparing the expression profiles within each pairs of sub-modules which was compared with 5000 surrogate maps accounting for spatial autocorrelation. The $p$ values based on the null distribution is shown with significance levels * $p < 0.05$ ** $p < 0.01$. B) Bold load effect (βBOLD) time series during a trial within each sub-module. The shaded area corresponds to standard error across regions within the respective module.

| Model 1 | BOLD ~ $\alpha$2a + 1 + ($\alpha$2a + 1 \| Subject) | | | | |
|---|---|---|---|---|---|
| | **Fixed effect** | | | Random effect | |
| | Beta ± $\sigma$ | t (5992) | *p* value | Beta | CI (95%) |
| $\beta_0$ | -0.023 ± 0.01 | -2.3753 | 0.017 | 0.034 | - |
| $\beta_{\alpha 2a}$ | 0.037± 0.016 | 2.29 | 0.022 | 0.056 | [0.053, 0.060] |
| Stats | AIC | BIC | LogLikelihood | Deviance | |
| | -11985 | -11944 | 5998.3 | -11997 | |

| Model 2 | PC ~ $\alpha$2a + 1 + ($\alpha$2a + 1 \| Subject) | | | | |
|---|---|---|---|---|---|
| | **Fixed effect** | | | Random effect | |
| | Beta ± $\sigma$ | t (5992) | *p* value | Beta | CI (95%) |
| $\beta_0$ | 0.004 ± 0.002 | 1.871 | 0.061 | 0.009 | [0.006, 0.013] |
| $\beta_{\alpha 2a}$ | -0.001±0.003 | -0.51 | 0.6 | 0.010186 | [0.006, 0.017] |
| Stats | AIC | BIC | LogLikelihood | Deviance | |
| | -31198 | -31158 | 15605 | -31210 | |

| Model 3 | PC ~ BOLD + 1 + (BOLD + 1 \| Subject) | | | | |
|---|---|---|---|---|---|
| | **Fixed effect** | | | Random effect | |
| | Beta ± $\sigma$ | t (5992) | *p* value | Beta | CI (95%) |
| $\beta_0$ | 0.003 ± 0.001 | 2.784 | 0.0053 | 0.005 | [0.003, 0.007] |
| $\beta_{BOLD}$ | 0.0259 ± 0.006 | 3.961 | $7.55 \times 10^{-5}$ | 0.025 | [0.017, 0.037] |
| Stats | AIC | BIC | LogLikelihood | Deviance | |
| | -31346 | -31306 | 15679 | -31358 | |

**Supplementary Table S1**: Mixed effect models relating $\alpha$2a receptors atlas, Bold parametric effect, parametric participation coefficient (PC). Then regressors for both the fixed effect and the random effect (with subjects as grouping factor) is shown with the respective t test and p value. A significant effect is shown in the fixed effect of model 1 and model 3. Random effects were significant in the three models.

**Supplementary table S2:** list of all 375 pre-defined regions-of-interest (ROI) and its module assignment. There are 333 cortical parcels (161 and 162 regions from the left and right hemispheres, respectively) using the Gordon atlas (4), 14 subcortical regions from Harvard-Oxford subcortical atlas (bilateral thalamus, caudate, putamen, ventral striatum, globus pallidus, amygdala and hippocampus; http://fsl.fmrib.ox.ac.uk/), and 28 cerebellar regions from the SUIT atlas (5). The table is grouped by module assignment (i.e. blue; red; green and yellow; Figure 2 of main text); the network they belong, and the hemisphere when possible to the left and right column, marked in L or R, respectively. The regions that where used in the circle plot from Figure 2B (main text) are shaded in dark grey. DAN, dorsal attention; VN, visual; FPN, frontoparietal; SN, salience; CO, cingulo-opercular; VAN, ventral attention; SMm, somatomotor mouth; SMh, somatomotor hand; RSpN, retrosplenial; FTPN, frontotemporal; DMN, default mode; AN, auditory; CPN, cinguloparietal; SubC, subcortex; Cer, Cerebellar.

| Parcel ID | Hem | Network | Region | Parcel ID | Hem | Community | Region | Module |
|---|---|---|---|---|---|---|---|---|
| 65 | L | AN | L_PI | 224 | R | AN | R_PIns | Blue |
| 67 | L | AN | L_STG | 233 | R | AN | R_STG | Blue |
| 69 | L | AN | L_STG | 181 | R | CON | R_mPMA | Blue |
| 104 | L | AN | L_LPCG | 185 | R | CON | R_rACC | Blue |
| 63 | L | CON | L_OPJ | 187 | R | CON | R_rACC | Blue |
| 76 | L | CON | L_Mins | 188 | R | CON | R_rACC | Blue |
| 82 | L | CON | L_Sins | 219 | R | CON | R_OPJ | Blue |
| 103 | L | CON | L_OPJ | 238 | R | CON | R_Sins | Blue |
| 105 | L | CON | L_OPJ | 245 | R | CON | R_Mins | Blue |
| 112 | L | CON | L_LPCG | 246 | R | CON | R_Mins | Blue |
| 93 | L | CPN | L_mOPJ | 274 | R | CON | R_sAI | Blue |
| 43 | L | DAN | L_FEF | 318 | R | CON | R_FP | Blue |
| 52 | L | DAN | L_SPL | 250 | R | DAN | R_IFG | Blue |
| 55 | L | DAN | L_LSPL | 220 | R | DMN | R_OPJ | Blue |
| 88 | L | DAN | L_SPL | 257 | R | DMN | R_sPCC | Blue |
| 92 | L | DAN | L_mSPL | 167 | R | FPN | R_SPL | Blue |
| 95 | L | DAN | L_pOPJ | 272 | R | FPN | R_DLPFC | Blue |
| 106 | L | DAN | L_lPMA | 280 | R | None | R_mPFC | Blue |
| 121 | L | None | L_OFC | 284 | R | None | R_mPFC | Blue |
| 122 | L | None | L_OFC | 285 | R | None | R_mPFC | Blue |
| 123 | L | None | L_OFC | 286 | R | None | R_mPFC | Blue |
| 124 | L | None | L_OFC | 287 | R | None | R_mPFC | Blue |
| 125 | L | None | L_OFC | 288 | R | None | R_mPFC | Blue |
| 35 | L | SMh | L_mS1 | 302 | R | None | R_vVis | Blue |
| 37 | L | SMh | L_mM1 | 303 | R | None | R_vVis | Blue |
| 57 | L | SMh | L_S1 | 306 | R | None | R_vVis | Blue |
| 58 | L | SMh | L_S1 | 195 | R | SMh | R_mM1 | Blue |
| 3 | L | SMm | L_lM1 | 164 | R | SMm | R_lM1 | Blue |
| 39 | L | SMm | L_lM1 | 197 | R | SMm | R_lM1 | Blue |
| 59 | L | SMm | L_S1 | 212 | R | SMm | R_lM1 | Blue |
| 83 | L | SN | L_Ains | 247 | R | SN | R_Ains | Blue |
| 61 | L | VAN | L_TOJ | 221 | R | VAN | R_TOJ | Blue |
| 75 | L | VAN | L_Sins | 241 | R | VAN | R_FO | Blue |
| 79 | L | VAN | L_FO | 242 | R | VAN | R_FO | Blue |
| 158 | L | VAN | L_PMA | 256 | R | VN | R_smVis | Blue |
| 90 | L | VN | L_smVis | 258 | R | VN | R_lVis | Blue |
| 97 | L | VN | L_lVis | 263 | R | VN | R_V1 | Blue |
| 98 | L | VN | L_lVis | 307 | R | VN | R_V1 | Blue |
| 139 | L | VN | L_lVis | 308 | R | VN | R_V1 | Blue |
| 141 | L | VN | L_V1 | 309 | R | VN | R_V1 | Blue |
| **Subcortical and cerebellar regions** | | | | | | | | |
| 353 | | Cer | Vermis_VI | 354 | | Cer | Right_VI | Blue |
| 356 | | Cer | Vermis_CrusI | 362 | | Cer | Vermis_VIIb | Blue |
| 361 | | Cer | Left_VIIb | 365 | | Cer | Vermis_VIIIa | Blue |

| | | | | | | | | |
|---|---|---|---|---|---|---|---|---|
| 370 | | Cer | Left_IX | 340 | | SubC | Left Accumbens | Blue |
| 374 | | Cer | Vermis_X | 342 | | SubC | Right Caudate | Blue |
| 337 | | SubC | Left Pallidum | 347 | | SubC | Right Accumbens | Blue |
| 335 | | SubC | Left Caudate | | | | | Blue |
| 64 | L | AN | L_OPJ | 230 | R | AN | R_OPJ | Red |
| 21 | L | CON | L_PCC | 180 | R | CON | R_mPMA | Red |
| 34 | L | CON | L_mPMA | 192 | R | CON | R_mPMA | Red |
| 40 | L | CON | L_dPMA | 196 | R | CON | R_mPMA | Red |
| 84 | L | CON | L_sAIns | 198 | R | CON | R_dPMA | Red |
| 41 | L | DAN | L_FEF | 223 | R | CON | R_OPJ | Red |
| 42 | L | DAN | L_FEF | 248 | R | CON | R_sAIns | Red |
| 49 | L | DAN | L_dPMA | 249 | R | CON | R_sAIns | Red |
| 51 | L | DAN | L_SPL | 189 | R | DAN | R_mSPL | Red |
| 87 | L | DAN | L_SPL | 199 | R | DAN | R_FEF | Red |
| 91 | L | DAN | L_mSPL | 203 | R | DAN | R_FEF | Red |
| 2 | L | SMh | L_mS1 | 208 | R | DAN | R_FEF | Red |
| 30 | L | SMh | L_mS1 | 211 | R | DAN | R_SPL | Red |
| 47 | L | SMh | L_M1 | 163 | R | SMh | R_mS1 | Red |
| 48 | L | SMh | L_PMA | 193 | R | SMh | R_mS1 | Red |
| 50 | L | SMh | L_S2 | 205 | R | SMh | R_PMA | Red |
| 54 | L | SMh | L_lS1 | 206 | R | SMh | R_PMA | Red |
| 56 | L | SMh | L_S2 | 207 | R | SMh | R_PMA | Red |
| 60 | L | VAN | L_TOJ | 209 | R | SMh | R_S1 | Red |
| 5 | L | VN | L_dVis | 210 | R | SMh | R_S1 | Red |
| 99 | L | VN | L_lVis | 213 | R | SMh | R_lS1 | Red |
| 138 | L | VN | L_iV1 | 214 | R | SMh | R_S1 | Red |
| | | | | 215 | R | SMh | R_S1 | Red |
| | | | | 217 | R | SMh | R_S1 | Red |
| | | | | 270 | R | SMh | R_lS1 | Red |
| | | | | 228 | R | VAN | R_TOJ | Red |
| | | | | 166 | R | VN | R_sVis | Red |
| | | | | 251 | R | VN | R_SPL | Red |
| | | | | 255 | R | VN | R_smVis | Red |
| | | | | 264 | R | VN | R_lV1 | Red |
| | | | | 267 | R | VN | R_lVis | Red |
| | | | | 252 | R | DAN | R_pSPL | Red |
| | | | | 262 | R | DAN | R_SPL | Red |
| | | | | 271 | R | DAN | R_dPMA | Red |
| | | | | 275 | R | DAN | R_dPMA | Red |
| **Subcortical and cerebellar regions** | | | | | | | | |
| 364 | | Cer | Left_VIIIa | 363 | | Cer | Right_VIIb | Red |
| 367 | | Cer | Left_VIIIb | 366 | | Cer | Right_VIIIa | Red |
| 373 | | Cer | Left_X | 369 | | Cer | Right_VIIIb | Red |
| 359 | | Cer | Vermis_CrusII | 375 | | Cer | Right_X | Red |
| | | | | 372 | | Cer | Right_IX | Red |
| 68 | L | AN | L_STG | 183 | R | SN | R_dACC | Green |
| 22 | L | CON | L_MCC | 317 | R | CON | R_SFG | Green |
| 147 | L | CON | L_FP | 254 | R | CPN | L_mOPJ | Green |
| 153 | L | CON | L_FP | 236 | R | DAN | R_IFG | Green |

| | | | | | | | | |
|---|---|---|---|---|---|---|---|---|
| 89 | L | CPN | L_mOPJ | 253 | R | DAN | R_pSPL | Green |
| 74 | L | DAN | L_FO | 266 | R | DAN | R_ITG | Green |
| 100 | L | DAN | L_ITG | 162 | R | DMN | R_PCC | Green |
| 107 | L | DAN | L_lPMA | 165 | R | DMN | R_SFG | Green |
| 110 | L | DAN | L_IFG | 200 | R | DMN | R_SFG | Green |
| 113 | L | DAN | L_lPMA | 225 | R | DMN | R_ITG | Green |
| 155 | L | DAN | L_DLPFC | 259 | R | DMN | R_OPJ | Green |
| 1 | L | DMN | L_PCC | 316 | R | DMN | R_SFG | Green |
| 4 | L | DMN | L_SFG | 325 | R | DMN | R_SFG | Green |
| 6 | L | DMN | L_OPJ | 326 | R | DMN | R_SFG | Green |
| 25 | L | DMN | L_pSMA | 168 | R | FPN | R_DLPFC | Green |
| 94 | L | DMN | L_LOFJ | 327 | R | FPN | R_SFG | Green |
| 114 | L | DMN | L_FP | 328 | R | FPN | R_DLPFC | Green |
| 126 | L | DMN | L_STS | 170 | R | FPN | R_ITG | Green |
| 154 | L | DMN | L_SFG | 182 | R | FPN | R_pSMA | Green |
| 156 | L | DMN | L_SFG | 240 | R | FPN | R_IFG | Green |
| 157 | L | DMN | L_SFG | 260 | R | FPN | R_SPL | Green |
| 7 | L | FPN | L_IFG | 261 | R | FPN | R_SPL | Green |
| 24 | L | FPN | L_rACC | 273 | R | FPN | R_DLPFC | Green |
| 78 | L | FPN | L_FP | 276 | R | FPN | R_dPMA | Green |
| 96 | L | FPN | L_SPL | 277 | R | FPN | R_FP | Green |
| 108 | L | FPN | L_DLPFC | 319 | R | FPN | R_FP | Green |
| 109 | L | FPN | L_DLPFC | 320 | R | FPN | R_FP | Green |
| 148 | L | FPN | L_FP | 281 | R | None | R_mPFC | Green |
| 149 | L | FPN | L_FP | 174 | R | RTN | R_RSp | Green |
| 115 | L | None | L_FP | 294 | R | RTN | R_vmVis | Green |
| 14 | L | RTN | L_RSp | 295 | R | RTN | R_vmVis | Green |
| 38 | L | SMh | L_S1 | 313 | R | RTN | R_HF | Green |
| 45 | L | SMh | L_M1 | 190 | R | SMh | R_mS1 | Green |
| 46 | L | SMh | L_M1 | 222 | R | VAN | R_TOJ | Green |
| 23 | L | VAN | L_pSMA | 229 | R | VAN | R_TOJ | Green |
| 62 | L | VAN | L_STS | 231 | R | VAN | R_FO | Green |
| 80 | L | VAN | L_iAI | 237 | R | VAN | R_FO | Green |
| 86 | L | VAN | L_FO | 333 | R | VAN | R_STS | Green |
| 8 | L | VN | L_vmVis | 169 | R | VN | R_vmVis | Green |
| 15 | L | VN | L_V1 | 175 | R | VN | R_V1 | Green |
| 16 | L | VN | L_V1 | 176 | R | VN | R_FFG | Green |
| 17 | L | VN | L_V1 | 177 | R | VN | R_FFG | Green |
| 20 | L | VN | L_RSp | 265 | R | VN | R_lV1 | Green |
| 131 | L | VN | L_vlVis | 293 | R | VN | R_vmVis | Green |
| 132 | L | VN | L_vlVis | 298 | R | VN | R_vVis | Green |
| 136 | L | VN | L_vlVis | 299 | R | VN | R_vVis | Green |
| 137 | L | VN | L_lV1 | 310 | R | VN | R_V1 | Green |
| 140 | L | VN | L_V1 | 311 | R | VN | R_V1 | Green |
| **Subcortical and cerebellar regions** | | | | | | | | |
| 348 | | Cer | Left_I_IV | 357 | | Cer | Right_CrusI | Green |
| 350 | | Cer | Left_V | 351 | | Cer | Right_V | Green |
| 355 | | Cer | Left_CrusI | 360 | | Cer | Right_CrusII | Green |
| 358 | | Cer | Left_CrusII | | | | | Green |
| 352 | | Cer | Left_VI | | | | | Green |
| 339 | | SubC | Left Amygdala | | | | | Green |

| | | | | | | | | |
|---|---|---|---|---|---|---|---|---|
| 10 | L | AN | L_PI | | | | | Yellow |
| 66 | L | AN | L_STG | 171 | R | AN | R_PIns | Yellow |
| 70 | L | AN | L_PIns | 227 | R | AN | R_STG | Yellow |
| 77 | L | AN | L_PIns | 232 | R | AN | R_STG | Yellow |
| 102 | L | AN | L_LPCG | 239 | R | AN | R_PIns | Yellow |
| 160 | L | AN | L_STS | 244 | R | AN | R_PIns | Yellow |
| 27 | L | CON | L_rACC | 268 | R | AN | R_LPCG | Yellow |
| 28 | L | CON | L_rACC | 269 | R | AN | R_LPCG | Yellow |
| 71 | L | CON | L_MIns | 329 | R | AN | R_PIns | Yellow |
| 72 | L | CON | L_MIns | 330 | R | AN | R_STG | Yellow |
| 81 | L | CON | L_Sins | 234 | R | CON | R_Mins | Yellow |
| 101 | L | CON | L_LPCG | 235 | R | CON | R_Mins | Yellow |
| 111 | L | CON | L_LPCG | 173 | R | CPN | R_PCC | Yellow |
| 12 | L | CPN | L_PCC | 184 | R | DMN | R_mPFC | Yellow |
| 26 | L | DMN | L_PCC | 186 | R | DMN | R_PCC | Yellow |
| 44 | L | DMN | L_SFG | 278 | R | DMN | R_FP | Yellow |
| 116 | L | DMN | L_mPFC | 279 | R | DMN | R_mPFC | Yellow |
| 117 | L | DMN | L_mPFC | 290 | R | DMN | R_STG | Yellow |
| 127 | L | DMN | L_STS | 315 | R | DMN | R_SFG | Yellow |
| 145 | L | DMN | L_SFG | 321 | R | DMN | R_SFG | Yellow |
| 146 | L | DMN | L_SFG | 322 | R | DMN | R_SFG | Yellow |
| 150 | L | DMN | L_mSFG | 323 | R | DMN | R_SFG | Yellow |
| 151 | L | DMN | L_SFG | 324 | R | DMN | R_SFG | Yellow |
| 152 | L | DMN | L_dACC | 331 | R | DMN | R_STG | Yellow |
| 9 | L | FPN | L_ITG | 226 | R | VAN | R_STS | Yellow |
| 85 | L | VAN | L_FO | 243 | R | VAN | R_iAI | Yellow |
| 161 | L | VAN | L_STS | 332 | R | VAN | R_STS | Yellow |
| 11 | L | None | L_HF | 172 | R | None | R_TempP | Yellow |
| 18 | L | None | L_FFG | 178 | R | None | R_FFG | Yellow |
| 19 | L | None | L_PCC | 179 | R | None | R_PCC | Yellow |
| 73 | L | None | L_IFG | 282 | R | None | R_mPFC | Yellow |
| 118 | L | None | L_ITG | 283 | R | None | R_mPFC | Yellow |
| 119 | L | None | L_OFC | 289 | R | None | R_STG | Yellow |
| 120 | L | None | L_OFC | 291 | R | None | R_STG | Yellow |
| 128 | L | None | L_TempP | 292 | R | None | R_TempP | Yellow |
| 129 | L | None | L_TempP | 296 | R | None | R_vVis | Yellow |
| 133 | L | None | L_vlVis | 297 | R | None | R_vVis | Yellow |
| 134 | L | None | L_TempP | 300 | R | None | R_vVis | Yellow |
| 135 | L | None | L_TempP | 301 | R | None | R_vVis | Yellow |
| 142 | L | None | L_PHG | 304 | R | None | R_vVis | Yellow |
| 144 | L | None | L_PHG | 305 | R | None | R_vVis | Yellow |
| 159 | L | None | L_STG | 312 | R | None | R_HF | Yellow |
| 13 | L | RTN | L_RSp | 314 | R | None | R_mTemp | Yellow |
| 130 | L | RTN | L_vlVis | 191 | R | SMh | R_mM1 | Yellow |
| 143 | L | RTN | L_PHG | 194 | R | SMh | R_mM1 | Yellow |
| 31 | L | SMh | L_mS1 | 201 | R | SMh | R_M1 | Yellow |
| 32 | L | SMh | L_mM1 | 202 | R | SMh | R_M1 | Yellow |
| 33 | L | SMh | L_mM1 | 204 | R | SMh | R_mM1 | Yellow |
| 36 | L | SMh | L_mM1 | 216 | R | SMh | R_S1 | Yellow |
| 53 | L | SMm | L_lM1 | 218 | R | SMm | R_lS1 | |
| 29 | L | SN | L_dACC | | | | | Yellow |
| **Sub cortical and cerebellar regions** | | | | | | | | |

| | | | | | | | | |
|---|---|---|---|---|---|---|---|---|
| 368 | | Cer | Vermis_VIIIb | 349 | | Cer | Right_I_IV | Yellow |
| 371 | | Cer | Vermis_IX | 341 | | SubC | Right Thalamus | Yellow |
| 334 | | SubC | Left Thalamus | 343 | | SubC | Right Putamen | Yellow |
| 336 | | SubC | Left Putamen | 344 | | SubC | Right Pallidum | Yellow |
| 338 | | SubC | Left Hippocampus | 345 | | SubC | Right Hippocampus | Yellow |
| | | | | 346 | | SubC | Right Amygdala | Yellow |
